# Learning predictive signals within a local recurrent circuit

**DOI:** 10.1101/2023.06.15.545081

**Authors:** Toshitake Asabuki, Colleen J. Gillon, Claudia Clopath

**Affiliations:** Department of Bioengineering, Imperial College London, London, UK

## Abstract

The predictive coding hypothesis proposes that top-down predictions are compared with incoming bottom-up sensory information, with prediction errors signaling the discrepancies between these inputs. While this hypothesis explains the presence of prediction errors, recent experimental studies suggest that prediction error signals can emerge within a local circuit, that is, from bottom-up sensory input alone. In this paper, we test whether local circuits alone can generate predictive signals by training a recurrent spiking network using local plasticity rules. Our network model replicates prediction errors resembling various experimental results, such as a biphasic pattern of prediction errors and context-specific representation of error signals. Our findings shed light on how synaptic plasticity can shape prediction errors and enables the acquisition and updating of an internal model of sensory input within a recurrent neural network.

## Introduction

The brain is thought to learn an internal model of the environment to predict upcoming sensory inputs (Keller & Mrsic-Flogel, 2018). In support of this hypothesis, a wide variety of experiments have reported mismatch responses in the brain (Garrido et al., 2009; Wacongne et al., 2012; Keller et al., 2012). These responses are typically elicited by presenting subjects with a series of familiar or consistent stimuli, and then introducing a deviant stimulus. Mismatch responses then emerge as the difference between the neural activity evoked by the standard stimulus and the deviant stimulus. For example, using electroencephalography (EEG), deviant stimuli have been shown to elicit a mismatch negativity (MMN) response in humans, typically seen as a biphasic response in which a negative deflection is followed by a positive one (Liebenthal et al., 2003; Luck 2005; Garrido et al., 2009; Wacongne et al., 2011; Nagai et al., 2017; Meyer et al., 2011). Early studies of this mismatch response were primarily conducted using auditory oddball paradigms (Liebenthal et al., 2003; Garrido et al., 2009). However, many studies have since shown MMN-like responses emerging from a variety of sensory tasks and brain regions, including visual (Strömmer et al., 2014; Kimura et al., 2009; Kimura et al., 2010; Meyer et al., 2011; Winkler et al., 2005) and auditory (Näätänen and Alho, 1997; Schröger and Wolff, 1996) areas, as well as cognitive processing regions like prefrontal cortex (Opitz et al., 2002; Näätänen et al., 2005). Although there may be some differences between these signals, together this suggests that mismatch responses may constitute a general mechanism for automatically detecting deviations from the brain’s internal model.

A plausible explanation for these mismatch signals in the brain is provided by the predictive coding hypothesis (Garrido et al., 2009). In predictive coding, at each level of sensory processing, top-down predictions from the brain’s internal model of the world are used to cancel out incoming sensory information. Only the discrepancies between the predicted and actual sensory information are communicated to higher levels of the cortical hierarchy. These discrepancies are called prediction errors and they are thought to be critical for the brain to improve its internal model, and thus the predictions passed down through the sensory processing hierarchy (Bastos et al., 2012; Rao & Ballard, 1999, Huang & Rao, 2011). Several generative models derived from the predictive coding hypothesis have been implemented in biologically plausible models of neurons to explain the mismatch signals observed in the brain. For example, Wacongne et al. (2012) proposed a spiking neuron model of predictive coding to account for the MMN in an oddball paradigm. Relatedly, a study by Lieder et al. (2013) based on dynamic causal models provided a potential link between the MMN and Bayesian filtering. In these models, mismatch responses are explained to be prediction errors, computed at different levels of the sensory processing hierarchy.

Although these types of generative predictive coding models provide a broad explanation of the emergence of mismatch signals, there are two important aspects that they do not capture. First, there is considerable evidence that the brain develops internal predictions through experience, and thus that learning and prediction errors occur alongside one another (Liebenthal et al., 2003; Wacongne et al., 2011; Fiser et al., 2016; Garrett et al., 2020; Hertäg & Clopath, 2022). However, it has not been demonstrated how a set of synaptic plasticity rules can account simultaneously for prediction errors and learning within the same network. Second, several studies have shown that mismatch responses can occur automatically in contexts during which there appears to be minimal top-down input, like when subjects are inattentive or even sleeping (Garrido et al., 2009). However, most existing models focus on the emergence of mismatch signals in hierarchical circuits, and do not explain how such signals can emerge within a local circuit, from bottom-up sensory input alone.

In this paper, we addressed these open questions by simulating a recurrent spiking network using local plasticity rules we recently proposed (Asabuki & Clopath, 2023). The network model consists of a population of excitatory and inhibitory spiking neuron models. The synapses onto excitatory neurons undergo synaptic plasticity, allowing them to develop connectivity patterns that predict the network activity evoked by upcoming sensory events. Simultaneously, inhibitory synapses undergo additional plasticity to maintain the excitatory-inhibitory balance. We found that recurrent networks trained with these plasticity rules replicated many features of prediction errors that have been observed in experimental studies (Liebenthal et al., 2003; Luck 2005; Garrido et al., 2009; Nagai et al., 2017; Wacongne et al., 2011; Meyer et al., 2011). For example, the prediction errors displayed a similar biphasic pattern to the MMN waveform, and were context-dependent (Audette & Schneider, 2023; Price et al., 2022). vOerall, our study provides new insights into the mechanism by which synaptic plasticity shapes prediction errors and the acquisition and updating of an internal model of sensory input within a recurrent neural network.

## Results

To test whether predictive signals can be computed in a local circuit, we simulated recurrent spiking network consists of excitatory (E) and inhibitory (I) model neurons (Fig. 1a). Excitatory neurons only were driven by external stimuli. We presented a number of stimuli to the network, each of which increased the firing rate of a non-overlapping subset of excitatory neurons. All feedforward connections were fixed.

**Figure 1.**
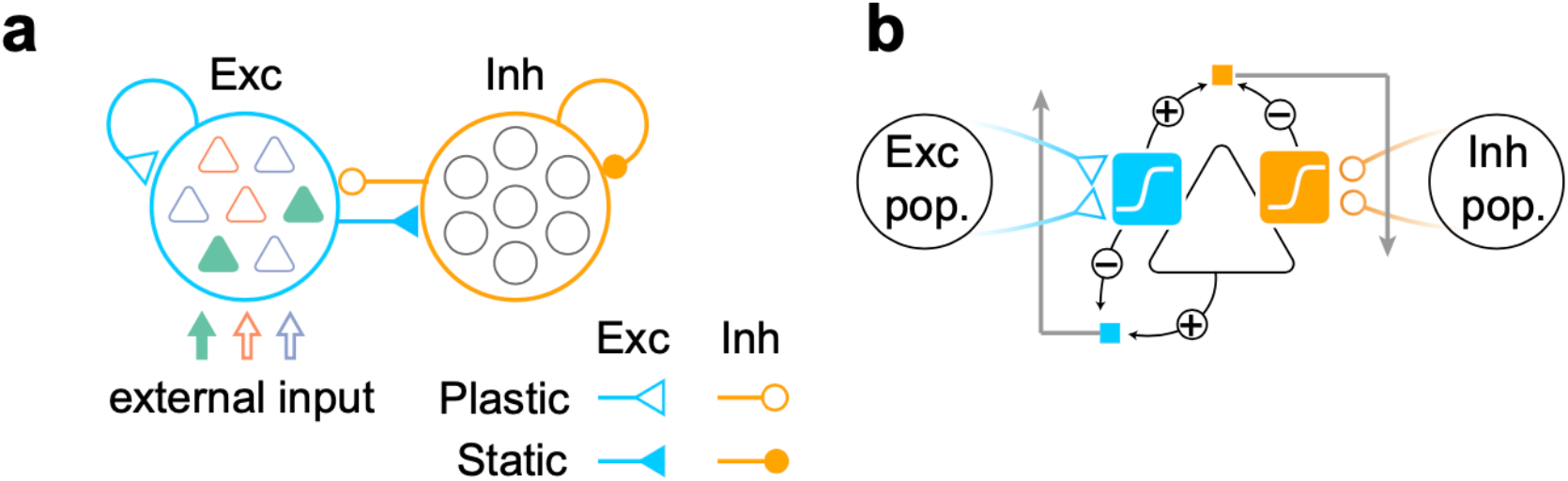
Model. (a) A network model with distinct excitatory and inhibitory populations. Only excitatory populations are driven by external inputs. Only synapses that project to excitatory neurons are assumed to be plastic. (b) A schematic of the plasticity rules proposed in (Asabuki & Clopath, 2023). Excitatory (blue) and inhibitory (orange) synapses projecting to an excitatory neuron (triangle) obey different plasticity rules. For excitatory synapses, errors between internally driven excitation (blue sigmoid) and the output of the cell provide feedback to the synapses and modulate plasticity (blue square). All excitatory connections seek to minimize such errors. For inhibitory synapses, the error between internally driven excitation (blue sigmoid) and inhibition (orange sigmoid) must be minimized to maintain excitation-inhibition balance (orange square).

We investigated how prediction errors are formed through sensory experiences by using synaptic plasticity rules that we proposed previously (Asabuki & Clopath, 2023). We assumed that excitatory and inhibitory synapses undergo distinct plasticity rules. Briefly, excitatory synapses that contributed to predicting neural activity were strengthened (Pfister et al., 2006; Urbanczik and Senn, 2014; Asabuki & Fukai, 2020; Asabuki & Fukai, 2023) (Fig. 1b, blue square), while the inhibitory synapses were modified to maintain the excitation-inhibition balance (EI balance) by predicting the recurrent excitatory potential (Fig. 1b, orange square).

### Emergence of a biphasic prediction error in a local recurrent network

Numerous experimental studies using electro-encephalography (EEG) and magneto-encephalography (MEG) have shown that a mismatch signal arises in auditory cortex when a rare “deviant” vocal stimulus occurs among a sequence of consistently repeated “standard” stimuli (Näätänen et al., 1978; Garrido et al., 2009; Strauss et al., 2015). This mismatch signal is measured by subtracting the response to the standard event from the response to the deviant one. Typically, this response difference (also known as difference waves) comprises both a negative (mismatch negativity; MMN) and a positive component (Liebenthal et al., 2003; Luck 2005; Garrido et al., 2009; Nagai et al., 2017; Wacongne et al., 2011; Meyer et al., 2011). Intriguingly, further experimental studies have found that inferotemporal cortex shows similar biphasic mismatch responses when a violation of transitional rules imposed during learning occurs (Meyer et al., 2011). Despite the consistency of these observations over various tasks, the plasticity mechanism that generates biphasic mismatch response is still unclear.

We first asked whether the proposed model could account for this biphasic mismatch response when a transition is violated in a learned sequence. To this end, a sequence with the deterministic transition “ABC” was presented to the network during a learning phase (Fig.2a, top). The excitatory synapses within each assembly (i.e., group of neurons targeted by the same stimulus, e.g. “A”) increased through learning, indicating the formation of cell assemblies for all stimuli (Fig.2b, diagonal blocks). Second, between-assembly connections for assembly A to B and B to C were strengthened, indicating that the model learned the transition probabilities between stimulus patterns as we have shown previously (Fig.2b, blue squares, Asabuki & Clopath, 2023).

We then investigated whether the network which learned stereotypical sequences showed a prediction error signal when a deviant sequence was presented. To this end, we measured the entire network response over a standard sequence (“ABC”) and a novel sequence with a deviant transition (“ABA”) (Fig.2a, bottom). In this analysis, all synaptic weights were fixed so that we could monitor the pure dynamics of the network. Note that the transition from pattern B to A in a deviant sequence violated the transition rule established during learning, and hence recurrent excitation connections from B to A had not been enhanced (Fig.2b, red square). In both the standard and deviant case, the network activities immediately after the transition showed an abrupt drop (early phase) followed by a slower rise (late phase) (Fig.2c, around vertical dashed line). However, we found a significant difference between the two: the response amplitudes were much stronger in both the negative and positive phases, in the deviant case than in the standard one, making the resulting error signals biphasic (Fig. 2d), consistent with results reported in the EEG literature (Liebenthal et al., 2003; Luck 2005; Garrido et al., 2009; Nagai et al., 2017; Wacongne et al., 2011; Meyer et al., 2011).

**Figure 2.**
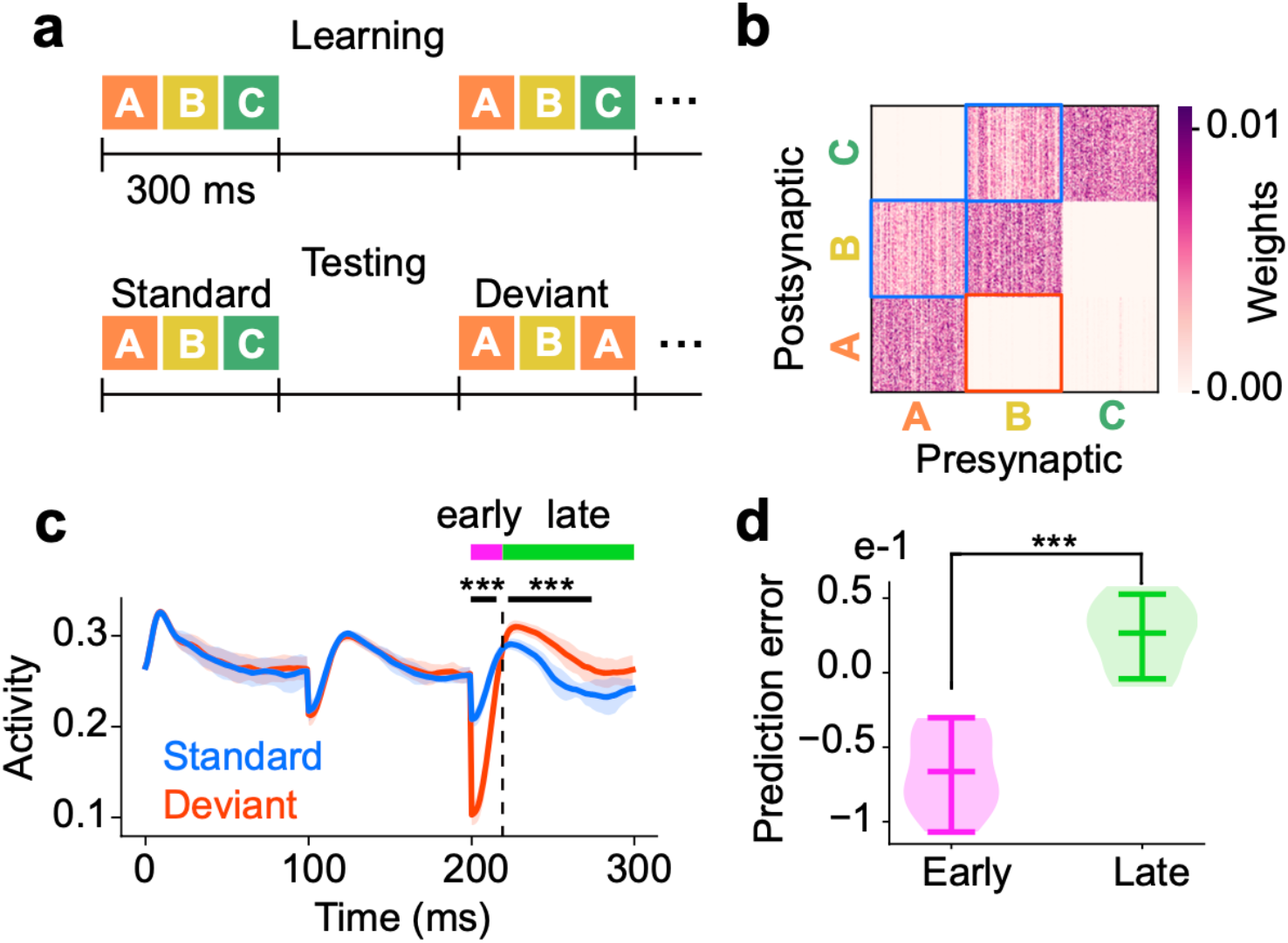
Biphasic prediction error learned through plasticity. (a, top) During learning, the sequence ‘ABC’ was repeatedly presented to the network. We assumed 300 ms-long gaps between each sequence. (a, bottom) After learning, all synapses were fixed and both the standard sequence ‘ABC’ and the deviant sequence ‘ABA’ were presented alternately. (b) Learned excitatory synapses are shown. Synapses were strengthened within each assembly (diagonal component of the matrix) and between assemblies that had transitions in a standard sequence (blue squares). Red squares show synapses between assemblies in the deviant sequence. (c) Mean responses of the whole network during the standard (blue) and deviant (red) sequences are shown. Period during which the last elements of sequences were presented was divided into early and late phase. Shaded areas represent s.d. over 10 trials. Black horizontal lines show periods during the two responses show significant difference. (d) Mean prediction errors during early and late phase over 10 independent simulations are shown. Here, prediction error was defined as difference between responses to deviant and standard sequences (deviant - standard). In c and d, p-values were calculated using a two-sided Welch’s t-test (***p<0.001).

In summary, these results show that our network model learns prediction error responses when presented with a stimulus sequence transition violation. In particular, the model shows a biphasic prediction error response, comprising a negative and a positive component, as found in neurophysiological experiments.

### Network mechanism of biphasic prediction errors

We then asked what is the network mechanism underlying this biphasic behavior. As the network dynamics were determined by recurrent connections onto assemblies, we analyzed the dynamics of excitatory and inhibitory recurrent currents. Here, we limited our analysis to the period during which the last stimulus of each sequence was presented, as the prediction error occurs only within this period. Specifically, we analyzed the currents in assembly C for the standard case and in assembly A for the deviant case (Supplementary Fig.1). We first explain the mechanism underlying the negative component of the prediction error, which occurs during the early phase. During the early phase in the standard sequence, excitatory and inhibitory currents showed almost similar levels, indicating that the model approximately maintained EI balance (Fig.3a, standard). In contrast, in the deviant case, these currents showed a significant deviation (Fig.3a, deviant; Fig.3b), breaking the EI balance. As the inhibitory current in the deviant case was dominant over the excitatory current, a negative component appeared in the prediction error in the early phase. Note that this break in EI balance was triggered by the significant decrease in excitatory currents in the deviant case (Fig.3a, cyan). Although the inhibitory current also showed similar behavior (Fig.3a, orange), the difference between the inhibitory currents was much smaller than for the excitatory currents (Supplementary Fig.3a). The significant difference in excitatory currents is likely explained by the fact that the recurrent connections from assembly B to A were not strengthened by learning, as we have already seen (Fig. 2b).

**Figure 3.**
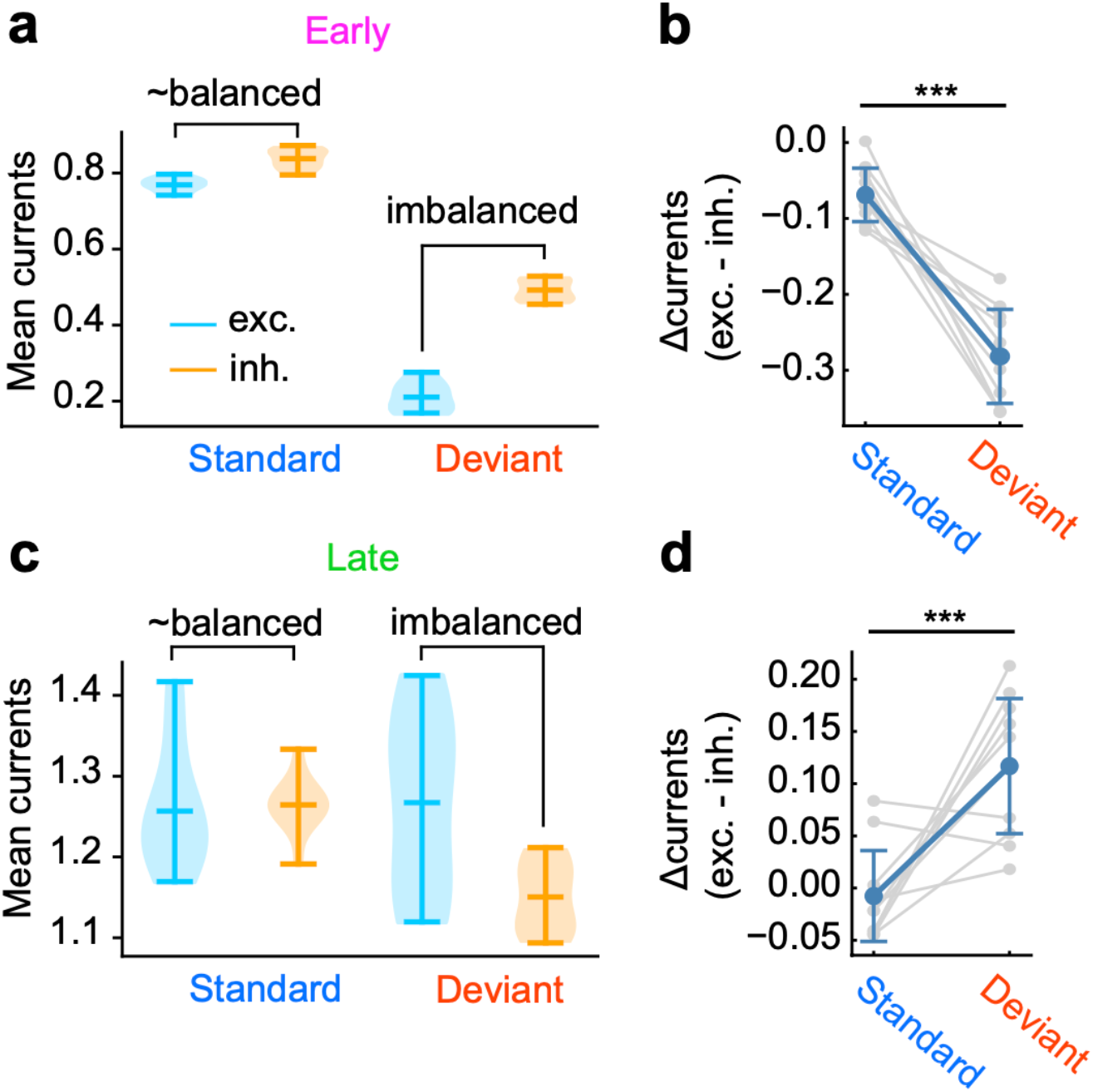
Analysis of recurrent currents. (a) Averaged excitatory (cyan) and inhibitory (orange) currents in the early phase for standard and deviant case are shown. The symbol “∼” indicates that an approximate EI balance is maintained in the standard case. (b) Differences between excitatory and inhibitory currents in the early phase for standard and deviant case are shown. (c) Same as a, but for the late phase. (d) Same as b, but for the late phase. In b and d, p-values were calculated using a two-sided Welch’s t-test (***p<0.001). Data points for each case were generated by 10 independent simulations.

The positive error component during the late phase is more surprising. We analyzed recurrent currents for both excitatory and inhibitory during the late phase, as we did for the early phase case. As in the early phase, the EI balance was maintained in the standard case (Fig.3c, standard) and was broken in the deviant case (Fig.3c, deviant; Fig.3d). Notably, we found that, in contrast to early phase, the excitatory current was much stronger than the inhibitory one in the deviant case, generating a positive prediction error during the late phase. The opposite direction of breaking EI balance was due to a significant decrease in inhibitory currents in the deviant case (Supplementary Fig.3b).

Altogether, these results suggest that negative and positive prediction errors in the early and late phases, respectively, result from distinct mechanisms. They show further that both negative and positive prediction errors can be explained by a disruption in the EI balance in the deviant case, but that the underlying mechanism is different for the two phases: in the early phase, the EI balance is broken due to the significant decrease in excitatory currents, whereas in the late phase, the break is due to a decrease in inhibitory currents.

### Learning of expectation dependent prediction error signals

The above results indicate that the model shows transitional surprise response if a predicted stimulus in a sequence is replaced by another stimulus, creating a deviant transition. If the network does indeed encode expected sequence order through learning, the responses to a same stimulus might be influenced by specific features of the stimuli that precede it.

Similarly, recent experimental study has shown that primary visual cortex (V1) responds differently to a stimulus based on whether the preceding stimulus in the sequence was expected or unexpected (Price et al., 2022). Extracellular neuronal recordings were acquired in awake head-fixed mice viewing sequences of visual stimuli “ABCD”, where each stimulus in the sequence had a set orientation. Mice were randomly assigned to four test days (i.e., days 1-4), such that each group experienced a different number of learning days (Fig. 4a). After experiencing the test stimuli once, mice were removed from the experiment. To quantify to what extent prediction certainty influences neural responses, two types of sequences (i.e., “AB*CD” and “EB*CD”) were used as test stimuli set in the experiment. Here, “E” was a novel or unexpected, stimulus which had not been included in the learned sequence “ABCD”. Further, “B*” reflects the fact that the “B” shown at test time varied in its orientation by a few degrees compared to the trained “B”. To summarize the experimental results, when comparing V1 neural activity mice tested on days 1 and 4 of learning, only the responses to stimulus “B*” in the “AB*” sequence, and not the “EB*” one, were significantly suppressed with learning.

**Figure 4.**
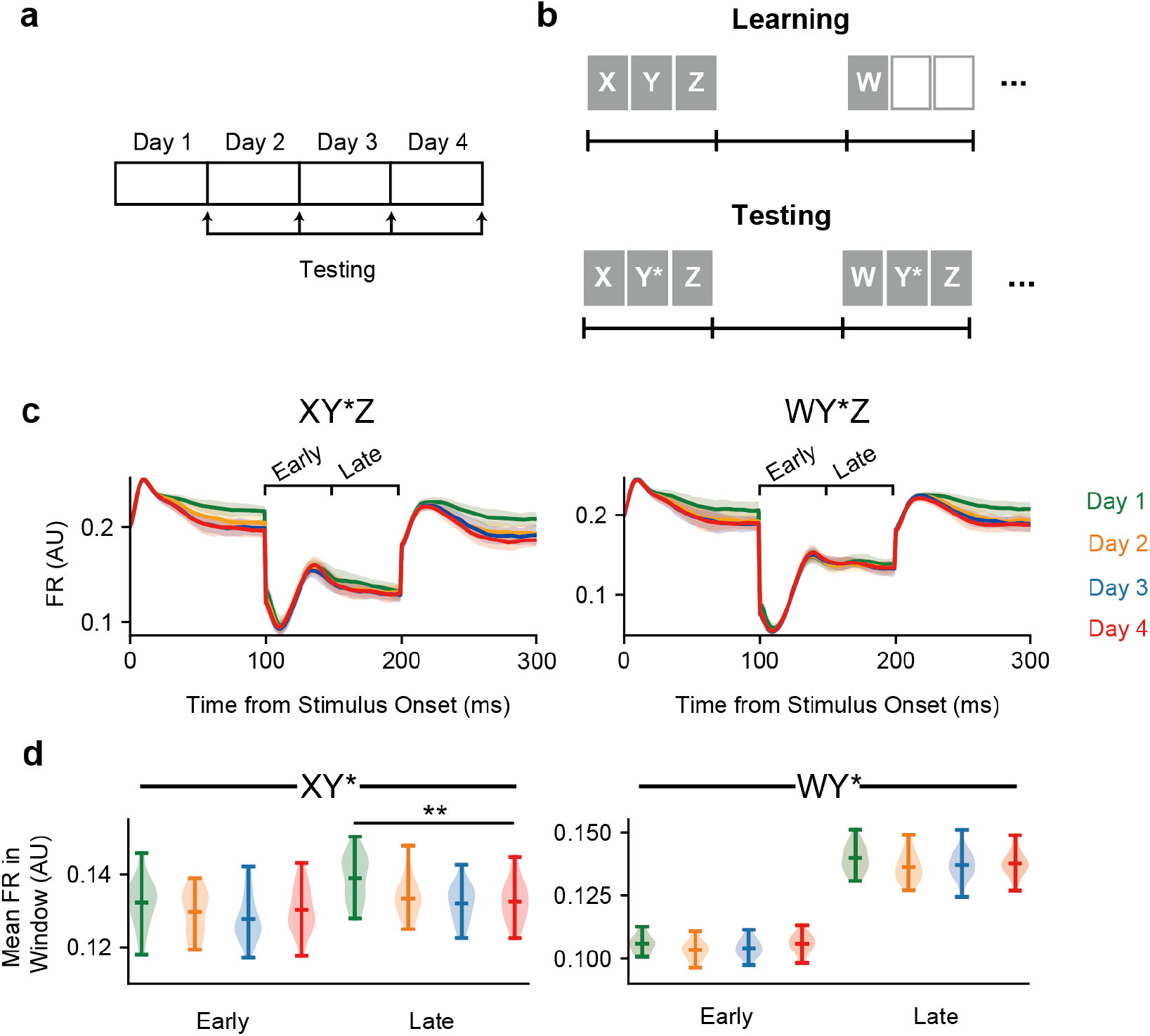
Learning of expectation dependent prediction error signals. (a) Learning phase was divided into four stages. At the end of each stage, all synapses were consolidated and network performance was tested against familiar and novel sequences. (b, top) During learning, sequence “XYZ” and an isolated element “W” were presented alternately. (b, bottom) During the testing phase, two sequences, “XY*Z” and “WY*Z”, were presented to the network alternately. Here, “Y*” is a noisy version of pattern “Y” (see Methods). (c, left) Mean network responses to the sequence “XY*Z” over different days. (c, right) Same as left figure, but for sequence “WY*Z”. (d) Average firing rates in the early and late windows after the occurrence of the second element “Y*” preceding “X” (left) or “W” (right) over different days are shown. Consistent with an experiment by Price et al. (2022), only the late window for “XY*” showed suppression through learning. In d, p-value was calculated using a two-sided Welch’s t-test (***p<0.001). Data points for each case were generated by 10 independent simulations.

We sought to test whether our model is consistent with these experimental results. In our simulation, a sequence “XYZ” and a single element sequence “W” were presented randomly during learning (Fig. 4b, top). Stimulus sequences were separated from each other by a 300 ms gap. As the learning progressed, recurrent weights formed four assemblies, with three of them (i.e., assemblies X, Y, and Z) being connected by unidirectional connections (as in Fig. 2, see Supplementary Fig. 3). We then asked how the maturation of weights influences prediction errors by comparing the response to two sequences (i.e., expected “XY*Z” and unexpected “WY*Z”), as in the experimental setting described above (Fig.4b, bottom). Here, during the presentation of “Y*”, we activated all assemblies, but stimulated assembly “Y” most strongly (Methods). To quantify to what extent these stimulation protocols influenced the network activity, we first split the period during which “Y*” was presented into an early and a late phase, as in the experiment (Fig. 4c). The average network activity for the expected and unexpected cases in the early and late phase was then calculated for each day. Consistent with experimental results, the network response to “Y*” was different between the pre- and post-learning stages, only for the “XY*” sequence (Fig.4d). A possible mechanism behind these results is that due to inhibitory synapses getting stronger with learning, the presentation of the expected sequence produced stronger inhibitory currents, leading to strong suppression in the expected case.

In summary, we have shown that the model network activity changes significantly through training, to a degree that is dependent on predictive certainty. Consistent with the experimental results, network responses to expected stimuli were suppressed in a late phase in a stimulus transition after training, but responses to novel stimuli did not show such suppression.

### Learning context-dependent prediction error signals

So far, we have explored how our network could develop a relatively simple class of prediction error-related activity. Indeed, in all the simulations so far, prediction errors were generated by a violation of a particular transition between a pair of stimuli. Although it is possible that prediction errors only encode a generic error signal, shared across multiple stimuli, a recent experimental study has shown that prediction error signals for particular stimulus emerges in a context-dependent manner (Audette & Schneider; 2023). The experiment showed that many neurons show strong suppression of responses to certain stimuli only in the expected context, indicating that these signals were not simply responses to the simultaneous presentation of multiple stimuli. As a simple mechanism in which shared inhibition among excitatory neurons does not explain such context-dependent signals based on expectation, this result suggests that inhibitory connections are precisely tuned to selectively generate expectations for sensory stimuli. However, the plasticity mechanism underlying learning such context-dependent prediction errors is still elusive. We therefore wondered whether our model could account for learning such context-dependent prediction error representations.

In Audette & Schneider (2023), mice were trained to perform a sound-generating spontaneous forelimb movement task to explore how movement-based predictions affect neural responses to expected and unexpected sounds. During training, a stereotypical auditory stimulus was presented at the beginning of each movement. After the animal underwent sufficient training to produce spontaneous forelimb movements, mice heard either the well-trained expected sound or a novel auditory stimulus with a slightly different frequency at the beginning of each lever movement they produced (‘active’ condition). These sounds were also played in conditions where the animal was not performing lever movements (‘passive’ condition). The experiment showed that neural responses to the expected sound in the active condition were suppressed compared to the same sound heard in the passive case. In contrast, responses to the unexpected sound under the active condition were enhanced relative to the passive case. This result suggests that prediction error signals emerge due to a specific combination of stimulus and context in a way that is dependent from expectation.

We show that our model can learn context-specific prediction errors as well. To this end, we considered two types of inputs: one corresponding to auditory signals, and the other to a motor signal from motor cortex. Excitatory populations were divided into two distinct populations, each of them receiving one auditory signals (i.e., familiar or novel; A or B) (Fig. 5a, top). All neurons, excitatory and inhibitory, received distributed step-shaped inputs to model motor command signals. We assumed that presentation of both auditory and motor command inputs increased excitatory drive to neurons targeted by each pattern. During learning, the auditory signals A and B were presented to the network alternately, with only signal A being combined to the motor signal (“active” condition, Fig. 5a, bottom).

**Figure 5.**
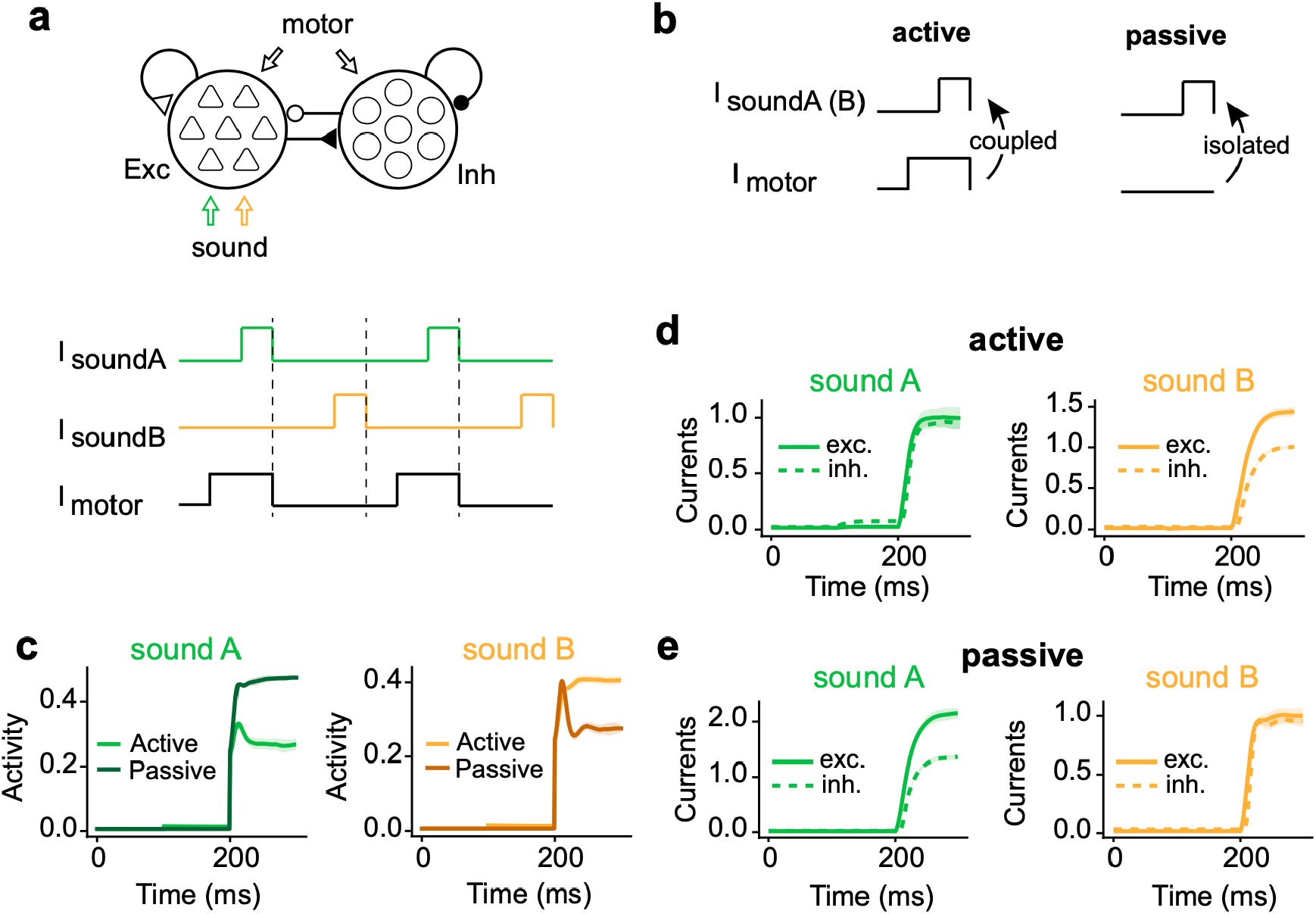
Learning of context-dependent prediction error signals. (a, top) Model settings. Excitatory population was divided to two subpopulations, each of which received either sound stimulus A or B (green and orange arrows). In addition, all neurons in the network could receive motor command input. (a, bottom) During learning, the auditory signal A was combined to the motor signal, but signal B was isolated from any motor signal. (b) After learning, sounds A and B were presented either coupled (active) or isolated (passive) from the motor input. (c) Mean network responses to sound A and B in the passive (Darker) and active (Lighter) context. (d) Recurrent excitatory and inhibitory currents while sound A (left) or B (right) was presented in the active context are shown. (e) Same as d, but in the passive context.

After learning, we first compared the network responses to the familiar stimulus (i.e., stimulus pattern ‘A’) under the active and the passive cases. We simulated responses under the active condition by measuring evoked dynamics in the network receiving both stimulus pattern A and a motor command signal (Fig. 5b, left). In contrast, we defined responses to stimulus ‘A’ under the passive condition as network responses to stimulus pattern ‘A’ alone (Fig. 5b, right). Consistent with the experimental results, the network responses to the expected sound in the active condition were suppressed compared to the responses to the same sound heard in the passive case (Fig. 5c, left). To test whether our model showed context-dependent prediction errors as shown experimentally (Audette & Schneider; 2023), we also compared the network responses to the stimulus pattern ‘B’ under the active and passive cases. Interestingly, in contrast to stimulus ‘A’, responses to the stimulus ‘B’ in the active condition were enhanced relative to the passive case (Fig. 5c, right).

We then asked what the potential mechanism is underlying the emergence of context-dependent prediction errors in the network model. As we have seen in the simple task, prediction error signals are generated by breaking the EI balance with an unexpected stimulus presentation. We therefore monitored excitatory and inhibitory recurrent currents during stimulus pattern ‘A’ or ‘B’, as presented under two different conditions (i.e., active or passive condition). When stimulus ‘A’ was presented, the strength of both currents was not drastically different in the active case (Fig. 5d, left), but was significantly different in the passive condition (Fig. 5e, left). In contrast, when the novel pattern ‘B’ was presented, the EI balance was maintained in the passive case (Fig. 5e, right), but broken in the active condition (Fig. 5d, right).

In summary, these results suggest that our network model with prediction-based plasticity can learn context-dependent prediction errors, as shown in the experimental study of Audette & Schneider (2023). The results also showed that prediction errors were generated due to a breaking in the EI balance that was precisely tuned in context-dependent manner.

## Discussion

In this study, we investigated how plasticity at recurrent excitatory and inhibitory synapses can produce prediction errors that carry features of mismatch responses observed in the brain. Specifically, we trained our model to predict upcoming network activity through its excitatory synapses, while its inhibitory synapses were tuned to maintain the EI balance (Asabuki & Clopath, 2023). We showed that the network learned the appropriate connectivity patterns to encode stimulus statistics and generate prediction error signals when deviant stimuli were presented, in agreement with various experimental results (Liebenthal et al., 2003; Luck 2005; Garrido et al., 2009; Nagai et al., 2017; Wacongne et al., 2011; Meyer et al., 2011; Audette & Schneider, 2023; Price et al., 2022).

Predictive coding suggests that mismatch signals may carry prediction errors in the brain. Indeed, previous computational studies have shown that spiking network models with layered cortical architecture trained using predictive coding replicate mismatch signals (Wacongne et al., 2012). Notably, however, although both predictive coding models and our recurrent network model generate prediction errors, there are several differences between the two. First, in standard predictive coding models, predicted state and prediction error signals are typically encoded in separate populations of neurons (Rao & Ballard, 1999; Bastos et al., 2012; Huang & Rao, 2011; Friston, 2012; Friston, 2018; Shipp, 2016). In contrast, in our model, both the predicted states and prediction errors are represented within a single neuron. We achieved this by implementing two distinct nonlinear activation functions within single neurons. A potential biologically plausible implementation of this feature would be to explicitly implement neurons as two-compartment units, where a dendritic compartment is non-linearly connected to a somatic compartment, which produces the neuron’s output. We leave the question of how exactly prediction and error signals might be encoded in more biologically plausible segregated neuron compartments and the associated plasticity rules to future work.

Another difference between standard predictive coding and our model is that, in predictive coding, top-down input predicts bottom-up input (Rao & Ballard, 1999; Bastos et al., 2012; Huang & Rao, 2011; Lotter et al., 2016). In contrast, in our model, predictions are generated locally, by the recurrent input. Experiments have shown that mismatch responses occur even when participants are engaged in a distractor task that draws attention away from the sensory modality in which the oddball stimulus occurs. This suggests that top-down input may not always be necessary for generating prediction error signals (Garrido et al., 2009; Näätänen et al., 2007).

We showed that local recurrent connectivity is sufficient to reproduce different forms of prediction error signals. Although a previous study suggested that combining recurrent connectivity with top-down prediction supports associative memory tasks via covariance learning, recurrent connections were trained to predict specifically in the spatial domain (Tang et al., 2023). Due to this nature of recurrent plasticity, the direct relationship between recurrent input and prediction error signals has remained elusive. We found that our model learns predictions in the temporal domain via local recurrent circuit and thus generates prediction error signals over sequential stimuli. How prediction error signals can encode both spatial and temporal information remains an open question.

Our model reproduces the general temporal profile of mismatch responses. Experimentally, several types of prediction error signals, including the standard MMN and inferotemporal cortical mismatch responses to visual stimuli violating sequence transitional rules, show a biphasic waveform (Nagai et al., 2017; Meyer et al., 2011). This waveform typically consists of an early negative deflection, followed by a positive one. Although one must be cautious in interpreting the meaning of positive and negative waves in EEG studies, it is notable that a very similar biphasic pattern emerges when measuring the difference in overall neural activity in response to standard and deviant stimuli in our recurrent network.

Predictive coding in which the prediction and prediction error signals are carried by different neurons (Rao & Ballard, 1999; Bastos et al., 2012; Huang & Rao, 2011; Friston, 2012; Friston, 2018), as opposed to a single neuron as in our study, may present some benefits. In particular, this may allow for more complex error signals to emerge through populations of error neurons specializing, for example, in positive and negative errors, respectively. Hertäg and Clopath (2022) show that such a network can, for example, use negative prediction error neurons to represent when an actual sensory stimulus is smaller than predicted, and positive prediction error neurons to represent when an actual sensory stimulus is bigger than predicted. In our model, it is conceivable that a similar function could be achieved using a temporal code, i.e., by modulating the amplitude of the positive or negative components of the prediction error signal, specifically. Indeed, we showed that both negative and positive prediction errors could arise from breaking of the EI balance. Specifically, negative prediction errors were due to a significant decrease in excitatory currents, whereas positive ones were due to a decrease in inhibitory currents. Further study is required to determine what advantages and disadvantages these different implementations present, and which best explains the spatial and temporal properties of mismatch signals in the brain.

Although our model can account for learning a variety of prediction error signals, in principle, it cannot learn predictions over the global structure of stimulus chunks. The reason behind this shortcoming is that our model learns local transition statistics between stimuli only, but is not designed to learn higher-order statistics (e.g., non-Markovian statistics) of stimuli. There are several possible ways to overcome this limitation. One possible solution is to consider much longer time scales than we considered in this study, such as calcium dynamics or NMDA spikes. Indeed, previous computational study show that NMDA-dependent plasticity is crucial for learning prediction error signals over global structure of sequence (Wacongne et al., 2012). Another possible way to achieve this is to consider hierarchically structured networks, as in real cortical structures. In such hierarchical networks, sub-networks lower in the hierarchy could learn to encode local element-level transitions, while the higher-level networks could learn slow and abstract dynamics (Maes et al., 2021), thus developing error signals related to the global structure of the sensory inputs. Extending our recurrent network model into a hierarchically structured model, and studying the relationship between recurrently driven and top-down driven prediction error signals could shed important light on the different between global and local mismatch signals in the brain.

In conclusion, our study sheds light on the learning mechanisms that may underlie mismatch signals in the brain, and provides a new perspective on the relationship between synaptic plasticity and prediction errors. Furthermore, it opens up new avenues for future studies of prediction error signals in hierarchical networks, and may contribute to the development of more flexible and biologically plausible models of neural computation.

## Methods

Our recurrent neural networks consist of *N*_*E*_ excitatory and *N*_*I*_ inhibitory neurons. During learning, the membrane potentials of neuron *i* at time *t* with external current 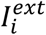 were calculated as follows:

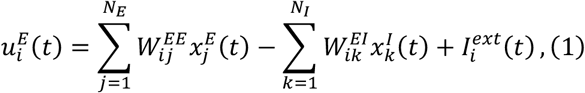

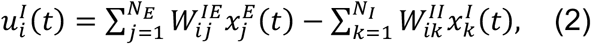

where 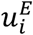 and 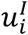 are the membrane potential of *i* -th excitatory and inhibitory neuron, respectively (see Table 1 for the list of variables and functions). The strength of external input 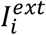 takes the value 1 if a stimulus pattern targeting neuron *i* was presented, and 0 otherwise. This structured external input was replaced by constant inputs 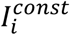 of value 0.3 during spontaneous activity. We will describe the details of stimulus patterns later. 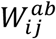 (*a, b* = *E; I*) is a recurrent connection weight from *j*-th neuron in population *b* to *i*-th neuron in population *a*. All neurons were connected with a coupling probability of *p =*0.5. The initial value of synaptic weights 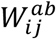 were uniformly set to 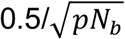. 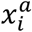 is a postsynaptic potential evoked by *i*-th neuron in population *a*, which will be described later.

**Table 1.** Definition of variables and functions.

|  |  |
| --- | --- |
| $u_i^E, u_i^I$ | Membrane potentials |
| $x_j^E, x_k^I$ | Postsynaptic potentials |
| $S_i^a$ | Poisson spike train generated by network neurons |
| $W_{ij}^{EE}, W_{ik}^{EI}, W_{ij}^{IE}, W_{ik}^{II}$ | Recurrent connections |
| $I_i^E, I_i^I$ | Synaptic currents generated by network neurons |
| $f_i^E, f_i^I$ | Instantaneous firing rates |
| $y_i^E, y_i^I$ | Recurrent predictions |
| $h_i$ | Memory trace |
| $\varphi$ | Sigmoidal function |

Spiking of each neuron model in population *E* was modeled as an inhomogeneous Poisson process with instantaneous firing rate 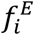 with a sigmoidal response function *φ*, with parameters slope *β* and threshold *θ*, as:

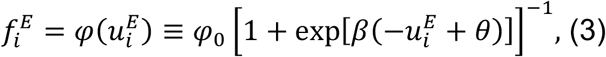

where *φ*_0_ is the maximum instantaneous firing rate of 50 Hz.

Inhibitory neurons’ firing rates were assumed to be calculated with static sigmoidal function as:

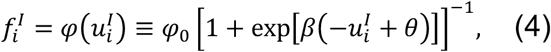

where the maximum instantaneous firing rate *φ*_0_ was assumed to be same as that of excitatory neurons (i.e., 50 Hz).

Neuron *i* in population *a* generates a Poisson spike train at the instantaneous firing rate of 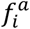. Let us describe the generated Poisson spike trains as:

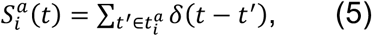

where *δ* is Dirac’s delta function and 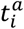 is the set of times at which a spike occurred in the neuron. The postsynaptic potential evoked by the neuron (i.e., 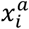) is then calculated as:

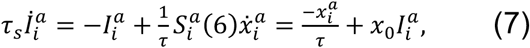

where *τ*_*s*_ *=* 5ms, *τ* = 15 ms, and *x*_0_ *=* 25.

### The learning rules

As in Asabuki & Clopath (2023), all excitatory synaptic connections onto excitatory neurons obeyed the following plasticity rule to predict the activity of postsynaptic neurons as:

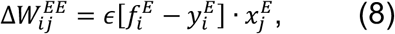

where 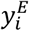 is a recurrent prediction of a firing rate, defined as:

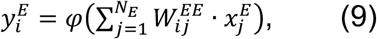

where the function *φ*(·) is the sigmoid function defined in Eq.3. In this study, the learning rate was set to *ϵ =* 10^−4^ in all simulations.

The inhibitory synapses onto excitatory neurons were plastic according to the following rule:

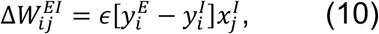

where 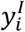 was the total inhibitory input onto postsynaptic neuron:

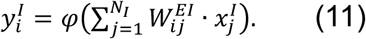

Through this inhibitory plasticity, inhibitory synapses were modified to maintain excitatory-inhibitory balance in all excitatory neurons.

### Simulation details

The parameters used in the simulations are summarized in Table 2. All simulations were performed in customized Python3 code written by TA with numpy 1.17.3 and scipy 0.18. Differential equations were numerically integrated using a Euler method with integration time steps of 1 ms.

**Table 2.**
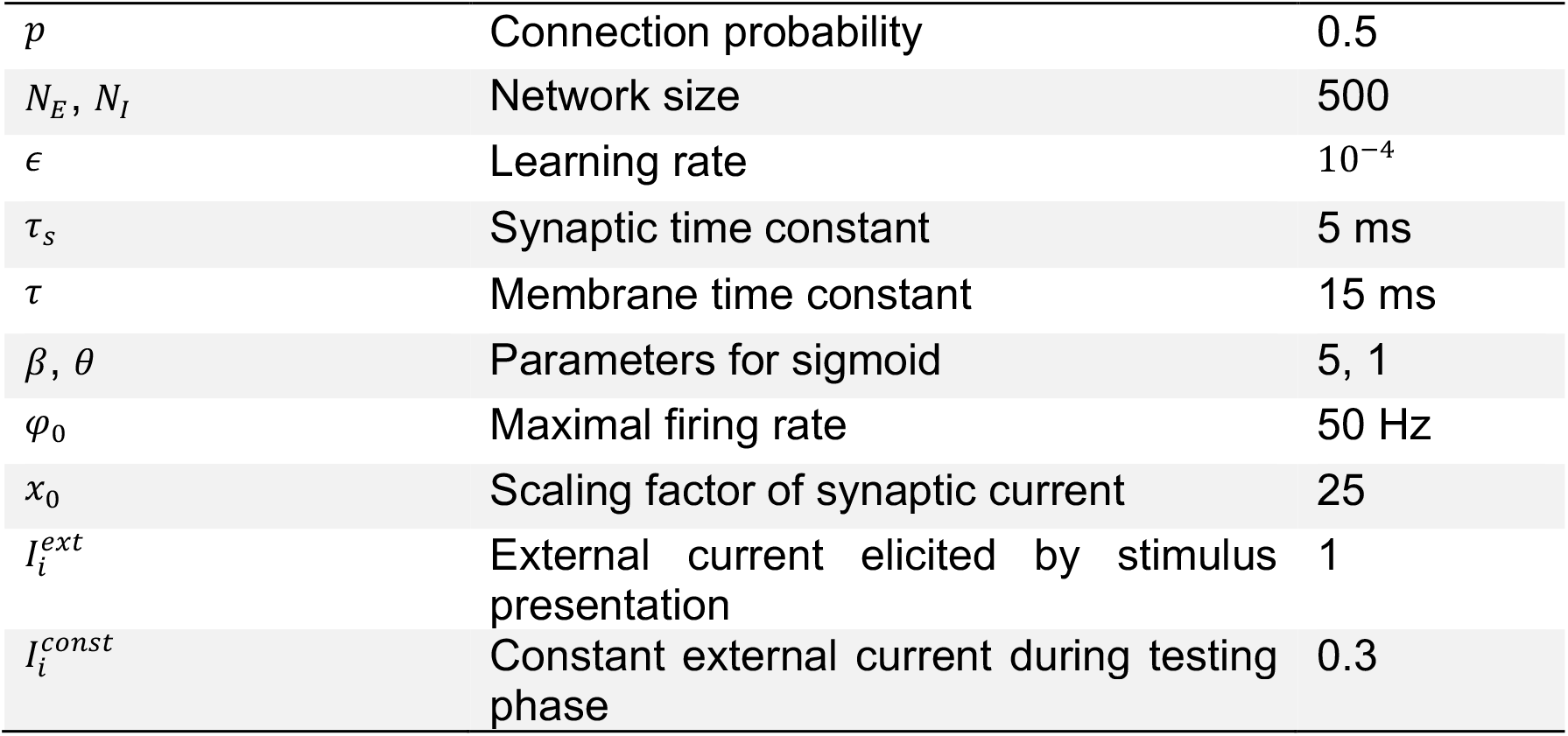
Parameter settings.

|  |  |  |
| --- | --- | --- |
| $p$ | Connection probability | 0.5 |
| $N_E, N_I$ | Network size | 500 |
| $\epsilon$ | Learning rate | $10^{-4}$ |
| $\tau_s$ | Synaptic time constant | 5 ms |
| $\tau$ | Membrane time constant | 15 ms |
| $\beta, \theta$ | Parameters for sigmoid | 5, 1 |
| $\varphi_0$ | Maximal firing rate | 50 Hz |
| $x_0$ | Scaling factor of synaptic current | 25 |
| $I_i^{ext}$ | External current elicited by stimulus presentation | 1 |
| $I_i^{const}$ | Constant external current during testing phase | 0.3 |

### Stimulation protocols

In all simulations, each stimulus pattern had a duration of 100 ms and was presented without an inter-pattern interval. Sequences of patterns were presented alternately with an interval of 300 ms. We assumed each neuron in a network was stimulated by one of the stimulus patterns and that targeted assemblies did not overlap. Presentation of each pattern triggered excitatory currents to its targeted neurons of strength 1 and zero otherwise. In all simulations, we assumed that only external inputs caused by the presentation of the stimuli were injected into the network during learning. In contrast, we assumed all excitatory neurons received both structured and constant background inputs over the whole period occurring after learning. During learning, all excitatory synaptic connections onto excitatory neurons were assumed to be plastic, while they were static during the testing phase after learning. The network was trained typically for 1,000 seconds except in Figure 4, where the simulation time was 300 seconds.

### Measuring prediction error signals

In Figure 2d, the early and late phases of responses were first defined as the periods before and after the point at which the mean standard and deviant responses intersected. Mean responses were calculated over 10 independent simulations. We then calculated the mean differences between responses over deviant and standard sequence (deviant-standard) for the two periods (i.e., early and late phases).

### Learning of expectation dependent prediction error signals

In Figures 4c and 4d, periods during which stimulus pattern “Y*” was presented were divided into a first (early) and a second (late) half. Mean activities over time were calculated over different days for each condition. To simulate stimulation with noisy stimulus “Y*”, assembly Y was stimulated with an input of intensity 0.6*I*^*ext*^ and the other assemblies with an input of intensity 0.4*I*^*ext*^, where *I*^*ext*^ is the strength of input for all other patterns (i.e., “X”, “W”, and “Z”).

## Acknowledgments

This work was supported by BBSRC (BB/N013956/1), Wellcome Trust (200790/Z/16/Z), the Simons Foundation (564408) and EPSRC(EP/R035806/1).

## Author Contributions

T.A. and C.C. conceived the study. T.A. performed the simulations and data analyses. And T.A., C.J.G. and C.C. wrote the paper.

