## Supplementary figures for "Learning predictive signals within a local recurrent circuit"

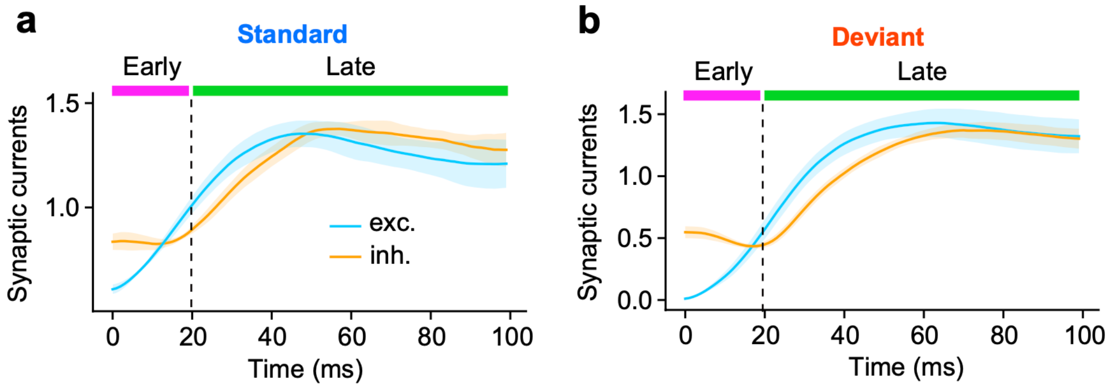


**Supplementary Figure 1.** (a) Excitatory (cyan) and inhibitory (orange) recurrent currents during the last element of standard sequence in Fig.2 was presented are shown. (b) Same with a, but for the deviant sequence.

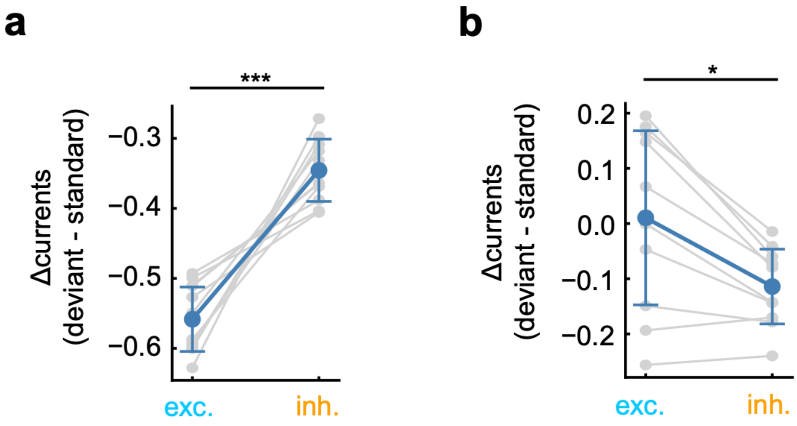


**Supplementary Figure 2.**(a) Differences in excitatory (cyan) or inhibitory (orange) currents between deviant and standard cases (deviant - standard) in the early phase are shown. (b)Same as a, but for late phase. In a and b, p-values were calculated using a two-sided Welch’ s t-test (*p<0.05, ***p<0.001). Data points for each case were generated by 10 independent simulations.

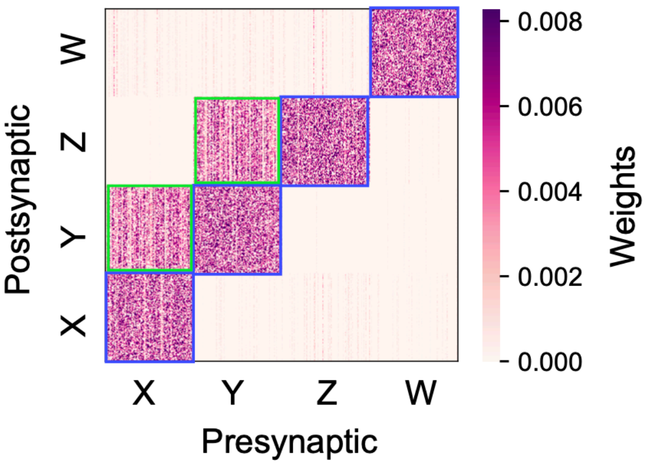


**Supplementary Figure 3.** Excitatory synapses learned to encode sequences shown in Figure 4b is shown. Synapses formed four assemblies (blue squares), with three of them (i.e., assemblies X, Y, and Z) being connected by unidirectional connections (grreen squares).
